# A Unified Neurocomputational Framework for Closed-Loop Motor Control and Sense of Agency

**DOI:** 10.64898/2026.09.11.750835

**Authors:** Johannes Nieuwenhuis, Silvestro Micera, Tommaso Bertoni

## Abstract

Motor control relies on the closed-loop comparison of motor commands and sensory feedback to correct errors and adapt to perturbations. Relevant features must be selected from a rich stream of sensory inputs and bound to the appropriate motor commands. This process remains poorly understood. Closed-loop control is accompanied by the experience of causing the observed feedback, sense of agency (SoA). SoA grounds self-identification, and its impairment is associated with lower prosthesis acceptance and disorders like autism and schizophrenia. Like closed-loop motor control, it relies on comparing desired and observed action outcomes in fronto-parietal circuits. Yet, these two phenomena have been studied independently, leaving SoA without a functional meaning and disregarding subjective aspects in motor control models. We propose that SoA is the subjective correlate of selecting self-caused sensory features for closed-loop control. We tested this in a visuomotor task where SoA was manipulated through temporal delays and adaptation to spatial perturbations served as a proxy for closed-loop integration. Delays similarly modulated SoA and closed-loop integration, suggesting these may emerge from the same phenomenon. We modelled our results in a Bayesian framework in which self-causation probability is inferred from temporal congruence, jointly modulating SoA and the weight attributed to visual feedback.

## Introduction

Precise motor control requires integrating sensory feedback and updating actions in a flexible manner. This process requires evaluating possible discrepancies between motor intentions and outcomes, and updating behaviour accordingly (Wolpert et al. 1998). To achieve this comparison in a precise and efficient manner, the selection of the relevant features from a rich stream of visual and somatosensory information, or sensory gating, is necessary (McGuire and Sabes 2009). Moreover, such feature selection for motor control co-occurs with the binding of those features with the relevant motor commands (McGuire and Sabes 2009). For example, during a precise manual task, motor commands associated with individual fingers need to be bound with the associated proprioceptive feedback to achieve effective control (Ghez et al. 1995). The principles regulating such gating and binding processes remain elusive.

Closed-loop motor control is normally accompanied by the subjective experience of being in control of our actions, also known as the sense of agency (Haggard and Chambon 2012). Like in models of motor control, the comparison between sensory feedback and motor efference copies is central in influential models of agency (Frith et al. 2000), which is broadly accepted to arise from the detection of a match between motor commands and reafferent sensory feedback. The subjective experience of agency over our bodily actions is rarely lost for most of the population, but crucial for the quality of life of specific groups of people. One such group is users of prosthetic devices, as the sense of agency over a prosthesis is a key predictor of its acceptance (Maimon-Mor and Makin 2020). Moreover, a reduced sense of agency is associated with psychiatric disorders such as schizophrenia and autism (Gallagher and Trigg 2016; Zalla and Sperduti 2015), and thought to contribute to shaping our sense of self as independent agents in the world (Gallagher 2000).

In addition to their functional similarity, closed-loop motor control and sense of agency show a significant neuroanatomical overlap. The posterior parietal cortex, a key area for integrating target and end effector position, passes on sensory information to the premotor cortex, to which it is connected bidirectionally (Archambault et al. 2015), specifically the intra-parietal sulcus. In turn, the cerebellum, as well as premotor and supplementary motor areas (SMA) send corrective signals to the primary motor cortex (M1). Similar fronto-parietal networks are also key for the sense of agency, with functional magnetic resonance studies converging on the dorsomedial prefrontal cortex, M1, pre-SMA, temporo-parietal junction and precuneus (Noel et al. 2025; Bertoni et al. 2025; Zito et al. 2020; Sperduti et al. 2011). Studies using non-invasive brain stimulation confirm the importance of such frontal (pre-SMA, (Cavazzana et al. 2015; Moore et al. 2010)) and parietal (angular gyrus, (Chambon et al. 2015)) regions for the sense of agency.

Notably, in spite of their functional and neuroanatomical common ground, studies on motor control and sense of agency have mostly constituted separate research lines. Despite its broad relevance, the sense of agency has not been conceptually linked to functionally relevant aspects of motor control, making it hard to objectively measure and operationalize. Previous works have hypothesized a link between visuomotor control and sense of agency (Reichenbach et al. 2014). Other works have suggested that a sense of agency may arise from the inference of self-causation (Legaspi and Toyoizumi 2019; Synofzik et al. 2009). However, these hypotheses have remained separated and still haven’t been tested experimentally. Conversely, despite playing a crucial role in motivation and reward (Karsh and Eitam 2015), and likely being related to lower-level aspects of motor control such as error correction (Wen et al 2021), sense of agency remains outside the scope of motor control models. Thus, the lack of an empirically supported theoretical framework accounting for both the sense of agency and closed-loop motor control constitutes a significant gap.

Here, we aim at unifying these domains, thus providing a functionally relevant account of the sense of agency, and a model of closed-loop motor control embedding the sense of agency as a relevant component. We propose that the sense of agency is the subjective counterpart of the selection of self-caused sensory features for closed-loop integration, and their binding with motor commands. Under this perspective, the brain would infer online which sensory features are generated by motor commands, based on sensorimotor congruences. Then, these features would be selectively integrated (bound) with the corresponding motor output to correct errors and allow precise motor control. On the subjective side, this process would be accompanied by the experience that our motor commands are the cause of those sensory features. Such self-causation inference would support the correction of small mismatches in actions that are overall perceived as self-caused, and sets the basis for learning new sensorimotor mappings. Unpredictable or extreme mismatches would instead yield a negative self-causation inference, leading to a loss of agency and gating sensory features out from closed loop control.

Being focused on the general principle of self-causation inference, our framework applies to any sensory modality integrated with motor commands (primarily proprioceptive and visual). For theoretical and practical reasons, we chose to empirically test our hypothesis by focusing on visuomotor integration as the most informative and general case. Practically, differently from proprioceptive feedback, visual feedback can be reliably manipulated experimentally, allowing us to precisely test our hypothesis within a quantitative framework. More importantly, visuomotor binding is particularly striking and complex, as humans are capable of binding motor commands with visual feedback coming from body parts, but also quickly learn new visuomotor mapping when using tools and external objects as effectors (Maravita and Iriki 2004; Wolpert et al. 2011). Such flexibility implies that visuomotor binding likely relies heavily on self-causation inference in order to identify the relevant effector and its mapping to motor commands. Crucially, visuomotor binding is effector-independent (Itaguchi and Fukuzawa 2014) and dissociable from visual attention (Reichenbach et al. 2014). Thus, visuomotor binding constitutes the ideal model for investigating the link between closed-loop motor control and sense of agency.

We thus developed a task that quantifies visuomotor integration during closed-loop control as the amount of adaptation to correctable perturbations in visual feedback. Simultaneously, we perturbed the sense of agency by introducing unpredictable temporal delays in visual feedback. We found that subjective ratings of agency and visuomotor integration were similarly affected by temporal delays, and co-varied even independently of the delay. This is in line with the idea that the sense of agency emerges from the process that gates information for closed-loop motor control. To formalize our findings, and corroborate the conceptual unification of agency and sensorimotor integration, we then showed that our results are well described by a Bayesian model where self-causation is inferred based on the congruence between motor commands and sensory feedback.

## Results

To investigate whether sense of agency is the subjective counterpart of feature selection for closed-loop integration, we developed a task allowing to independently measure sense of agency and the amount of visuomotor integration. Participants were instructed to perform circular movements with a joystick, while being presented with visual feedback about their motor behaviour through a small circular cursor on a screen (Figure 1A). After nine seconds of movement, they were asked to report their level of agency over cursor movements. Participants were explicitly instructed to only make circles with their hand, not with the cursor. We introduced an unpredictable, continuously varying temporal incongruence (from here, incongruence) to manipulate the subjects’ sense of agency, ranging from almost perfect congruence to an average delay of 0.6 seconds (see methods). In parallel, we presented correctable, predictable spatial distortions (from here, distortion) of the visual feedback, by modulating the gain of the response on either the x or y axis in a constant manner within a single trial (Figure 1B, 1C). The amount of adaptation to these spatial distortions was taken as a proxy of visuomotor integration, which we will refer to as the compensation index (CI, see methods and Figure 1C). A value of 0 indicates no visuomotor adaptation: visual feedback was ignored, and a perfect circle was performed with the joystick. A value of 1 indicates complete visuomotor adaptation: the distorted visual feedback was fully integrated in motor behaviour to produce a circular trajectory on the screen. To determine the optimal temporal window of analysis, we first evaluated from what point onwards stable adaptation was achieved within a single trial. Across all conditions, adaptation became consistent and reached a stable value from the fourth ellipse (Figure S1), out of an average of 16.7 (± 2.83 SD) ellipses per trial. We therefore used all ellipses from the fourth one onwards to calculate the CI for each incongruence.

**Figure 1.**
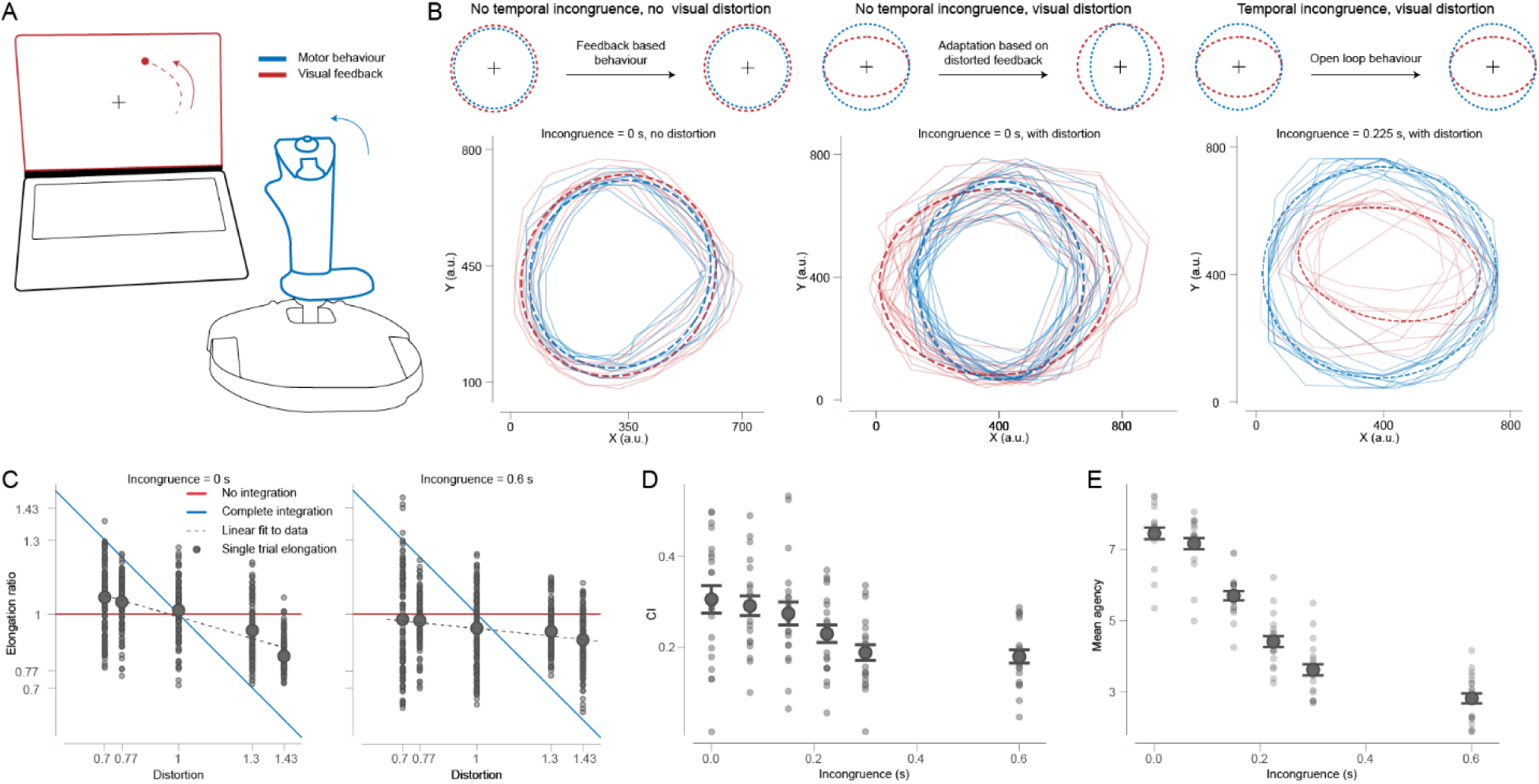
Behavioural task and main results. (A) Experimental set up. Participants were instructed to make circular movements with a hand-held joystick, while being presented with visual feedback on a screen. (B) Illustration of conditions in the experiment from three actual exemplary trials. We either presented no temporal incongruence and no spatial distortions (left), no temporal incongruence and spatial distortion (middle), temporal incongruence with spatial distortion (right) or without spatial distortion (not shown). In congruent trials, we expect participants to start with circular movements, and then either continue with circular movements in case of no distortion (left) or compensate for the distortion, by adjusting their movement in the opposite direction to the distortion (middle). In incongruent trials, we expect them to ignore the distortion and simply perform circular movements regardless of the distorted (right) or undistorted (not shown) feedback. (C) Extraction of the compensation index (CI) and example behaviour for congruent (left) and incongruent (right) trials. First, we fitted an ellipse to each individual elliptical joystick movement in a trial, and extracted its vertical elongation. If a participant simply performed circular movements irrespective of visual feedback, we expect the elongation to be independent of distortion (red line). If a participant fully compensated for the distortion, so that a perfect circle was drawn on the screen, we expect the elongation to be inversely proportional to the distortion (blue line). Thus, the CI is defined as the negative slope of the linear fit elongation∼distortion (dashed line). Larger circled dots indicate means per incongruence. (D,E) Effect of incongruence on the compensation index (D) and on agency ratings (E). Each dot indicates a participant, error bars indicate standard errors.

### Temporal incongruences similarly modulate visuomotor integration and sense of agency

Our framework predicts that temporal incongruence should lead to a negative self-causation inference, and thus to a decrease in the reported sense of agency for cursor movements. Conversely, this should gate visual feedback out of the control loop, leading to decreased CI values. This is qualitatively evident in Figure 1D, and was confirmed by a one-way ANOVA revealing a main effect of incongruence on the CI (generalized eta squared (GES) = 0.199, F(5,100) = 17.65, p<0.0001). Subjective agency ratings showed a similar downward trend with incongruence, also confirmed by an ANOVA (GES = 0.868 F(5,100) = 255.98, p<0.0001, Figure 1E). Subject-specific plots are shown in Figure S2 and S3. Importantly, according to our hypotheses and intended design, the correctable spatial distortions should have induced motor adaptation, but no strong modulation of agency compared to our temporal manipulation. This would justify their use as a proxy of visuomotor integration without affecting our key subjective outcome measure. A one-way ANOVA did reveal a significant effect of distortion on agency ratings (GES = 0.044, F(4,80) = 3.59, p = 0.01, Figure 2A). However, the effect size (GES) was much smaller than the effect of the temporal manipulation. Indeed, agency ratings averaged by incongruence exhibit a much broader (13.6 times) range of variability (7.45 ± 0.16 for incongruence = 0s to 2.82 ± 0.14 for incongruence = 0.6s) than when averaged by distortion (5.43 ± 0.11 for distortion = 1 to 5.09 ± 0.11 for distortion = 0.77). For all practical purposes, we can thus consider agency modulations to be dominated by incongruence.

**Figure 2.**
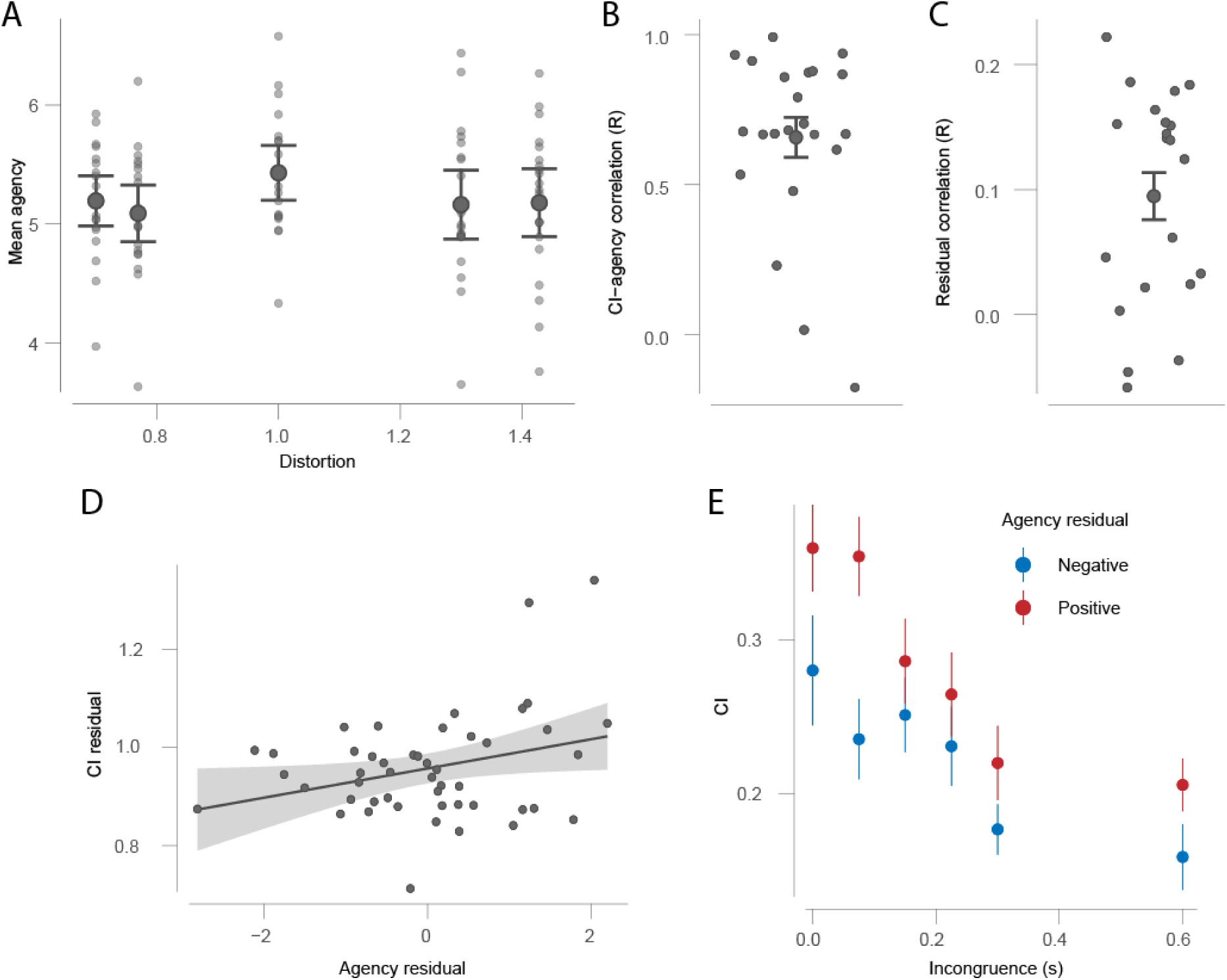
Relation between visuomotor integration and agency ratings. (A) Effect of distortion on agency ratings. Each dot indicates a participant, error bars indicate standard errors. (B) Correlation values between CI and agency ratings (averaged per incongruence) for each participant. Error bars represent standard errors. (C) Correlation between residual compensation and agency ratings, after regressing out the effect of temporal incongruence. Each dot indicates a participant, error bars represent standard errors. (D) Correlation between residual compensation and agency ratings for an example participant. Shades indicate 95% confidence intervals on the regression line. (E) CI values averaged across participants, split based on whether agency ratings were higher (red) or lower (blue) than average values for a given participant and incongruence (positive or negative residual agency). Error bars represent standard errors.

### Sense of agency and visuomotor integration co-vary independently of incongruence

We then directly investigated the link between visuomotor integration and sense of agency, by computing correlation values between mean agency ratings and CI for each participant. On average, CI values were highly correlated with agency ratings (R = 0.66 ± 0.14, Figure 2B), with only 4 out of 21 participants showing an R value below 0.5. This indicates that the compensation index is a good proxy of explicitly reported sense of agency. However, such correlation does not rule out that visuomotor integration and sense of agency may be two independent by-products of temporal incongruence. We thus investigated whether the CI and agency ratings would correlate even at fixed incongruence, which should be the case if they emerge from the same process rather than simply co-occurring. We subtracted to agency ratings and CI their average value per incongruence (see methods), obtaining residual values unaffected by incongruence. Indeed, 17 out of 21 subjects showed a positive correlation between CI and agency ratings, with correlation values being on average significantly positive (t(20) = 5.05, p < 0.0001, R = 0.105 ± 0.02, Figure 2C). In Figure 2D, we show an example subject in which the correlation between CI and agency is significant also at the single subject level. To visualize this result more intuitively, we plotted average CI values separately for trials in which subjects reported more or less agency than their individual average for each incongruence level (Figure 2E). This shows that participants tended to integrate visual feedback more in their motor behaviour when they had a comparatively stronger feeling of being in control of it, even at constant temporal incongruence.

### Incongruence modulates visuomotor integration and sense of agency independently of attention

In line with our framework and with previous literature (Reichenbach 2014), we expected the compensation index and the sense of agency to be independent of visual attention. To test this, we performed an additional experiment with a modified version of our task including only two incongruences (0 and 0.6 s), where we also measured the reaction time of participants to changes in cursor colour and size while performing the task. This was taken as a proxy of visual attention to the cursor. In half the trials, the dot turned red between six and nine seconds after trial start (sampled uniformly), to which participants were required to react as fast as possible by pressing the back trigger of the joystick. Our previous behavioural results were replicated in this modified version of the task. Figure 3A shows that the CI significantly decreased with incongruence (t(20) = 7.89, p<0.0001). The same was observed for the relation between agency and incongruence (t(20) = 12.71, p<0.0001, Figure 3B). Importantly, reaction times were not influenced by incongruence (Figure 3C), as confirmed by a paired t-test (t(20) = –0.43, p = 0.67). Additionally, the CI and mean agency ratings were not significantly correlated with reaction times in the congruent nor incongruent conditions. (Figure 3D-E, congruent agency R = 0.11 ± 0.42, p = 0.64, incongruent agency R = 0.00 ± 0.43, p=0.99, congruent CI R=0.03 ± 0.43, p=0.91, incongruent CI, R=0.03 ± 0.45, p=0.90; see also Figure S4).

**Figure 3.**
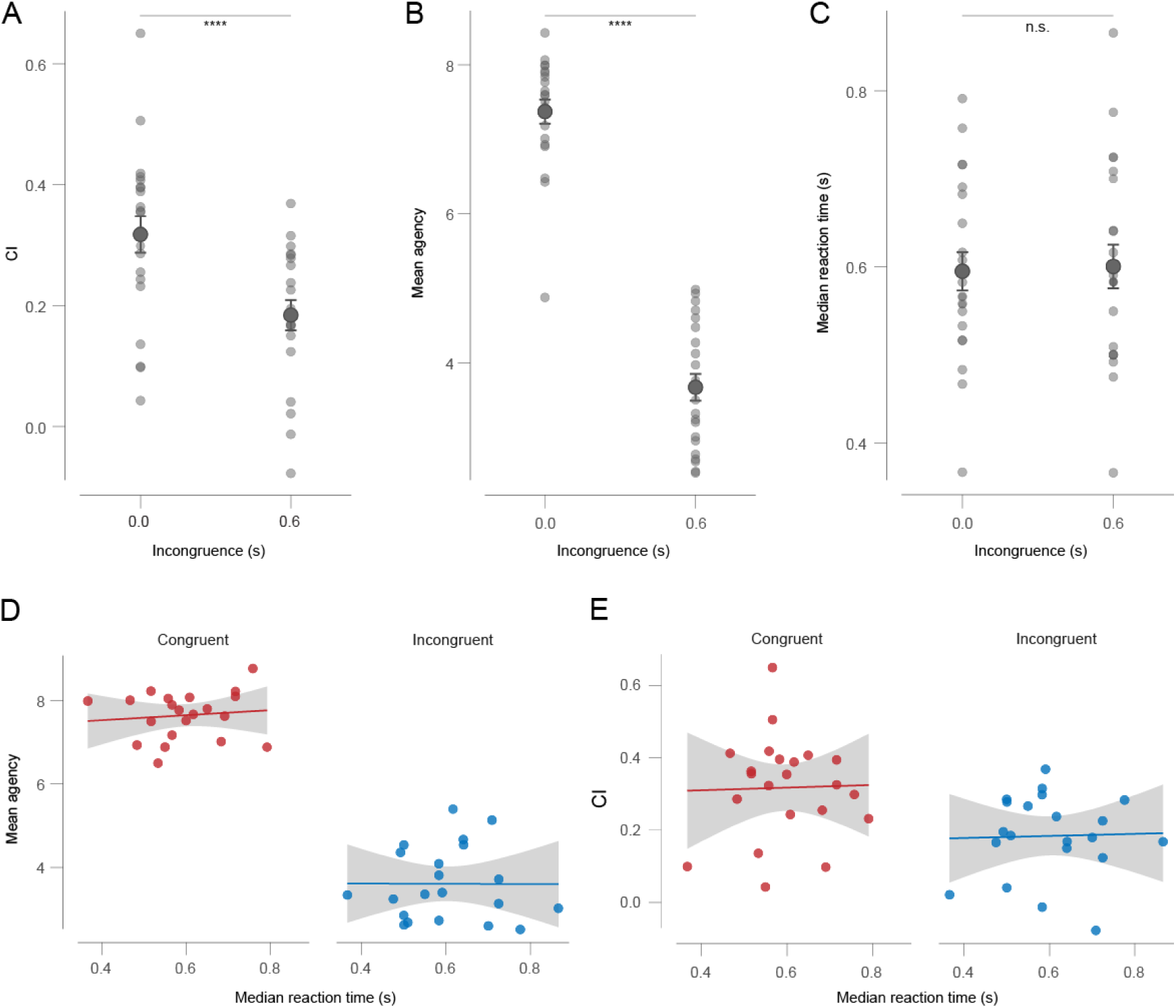
Relation between visual attention and incongruence. Panels (A) and (B) show respectively average CI and agency values for each participant in the congruent and incongruent conditions. Error bars represent standard errors (**** indicates p<0.0001). (C) Average reaction times to changes in cursor colour for congruent and incongruent trials. (D) Linear fit of mean agency ratings and median reaction time for congruent and incongruent condition. The solid line indicates the best fit, shades indicate 95% confidence intervals on the regression line. (E) Linear fit of mean CI and median reaction time for congruent and incongruent condition. The solid line indicates the best fit, shades indicate 95% confidence intervals on the regression line.

### Visuomotor integration is well modelled as the result of Bayesian self-causation inference

The similar functional structure of visuomotor integration and sense of agency, their shared neural substrate, and their empirically verified covariance begs the question: are sense of agency and visuomotor integration the subjective and functional counterparts of a shared substrate, the inference of self-causation?

To empirically test such an idea, and integrate it in a functional model of motor control, we developed a model describing motor behaviour as the result of a Bayesian causal inference process. In our proposed model, a subject will integrate more or less visual information in their motor commands depending on the inferred self-causation probability. If visual information about the dot’s trajectory is inferred to be self-caused, it is integrated with proprioceptive information to produce the desired motor behaviour, yielding high adaptation and thus a high CI. If visual information is inferred to be non-self-caused, it is disregarded and motor behaviour is adjusted based on proprioceptive inputs only, yielding low or no visuomotor adaptation/CI (Equation 1).

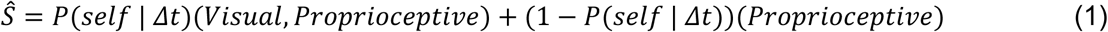

Where the motor behaviour *ŝ*, representing the elongation of the circular trajectories, arises from the weighted sum of fully closed loop behaviour based on the integration of visual and proprioceptive feedback (*Visual*, *Proprioceptive*) and semi open loop behaviour based on proprioceptive feedback only (from here, simply open loop). The relative weight of closed loop and open loop behaviour is determined by *P*(*self* | *Δt*), which is the inferred probability of being the cause of visual feedback given the temporal incongruence *Δt*. The exact relationship between CI and *P*(*self* | *Δt*) depends on the relative precisions of the visual and sensory modalities (see methods). However, for a given participant, the CI is always linearly proportional to *P*(*self* | *Δt*):

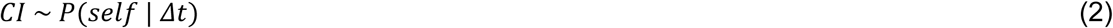

For an individual subject, *P*(*self* | *Δt*) is given by Equation 3:

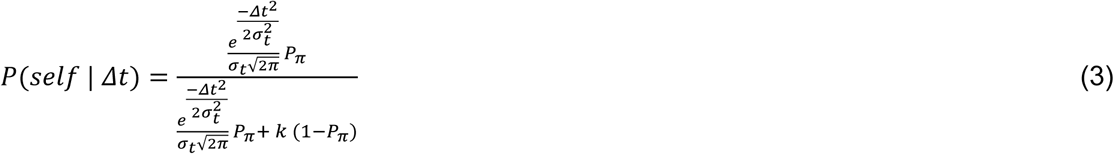

Where *σ_t_* is the sensory temporal uncertainty due to internal neural noise, *k* is a constant to represent the probability of non-self-causation given the delay and *P_π_* is the prior reflecting the overall tendency to attribute sensory feedback to the self. All terms in the model and the derivation of the equations are explained in depth in the methods section. For typical values of the parameters, *P*(*self* | *Δt*) starts at values close to 1 for 0 incongruence, and sharply drops towards 0 as *Δt* increases above the individual temporal precision *σ_t_* (Figure 4A). We optimized the free parameters of the model separately for every participant to best match individual behaviour (see methods). This allowed us to compute the CI predicted by the model, and compare it to measured behaviour.

**Figure 4.**
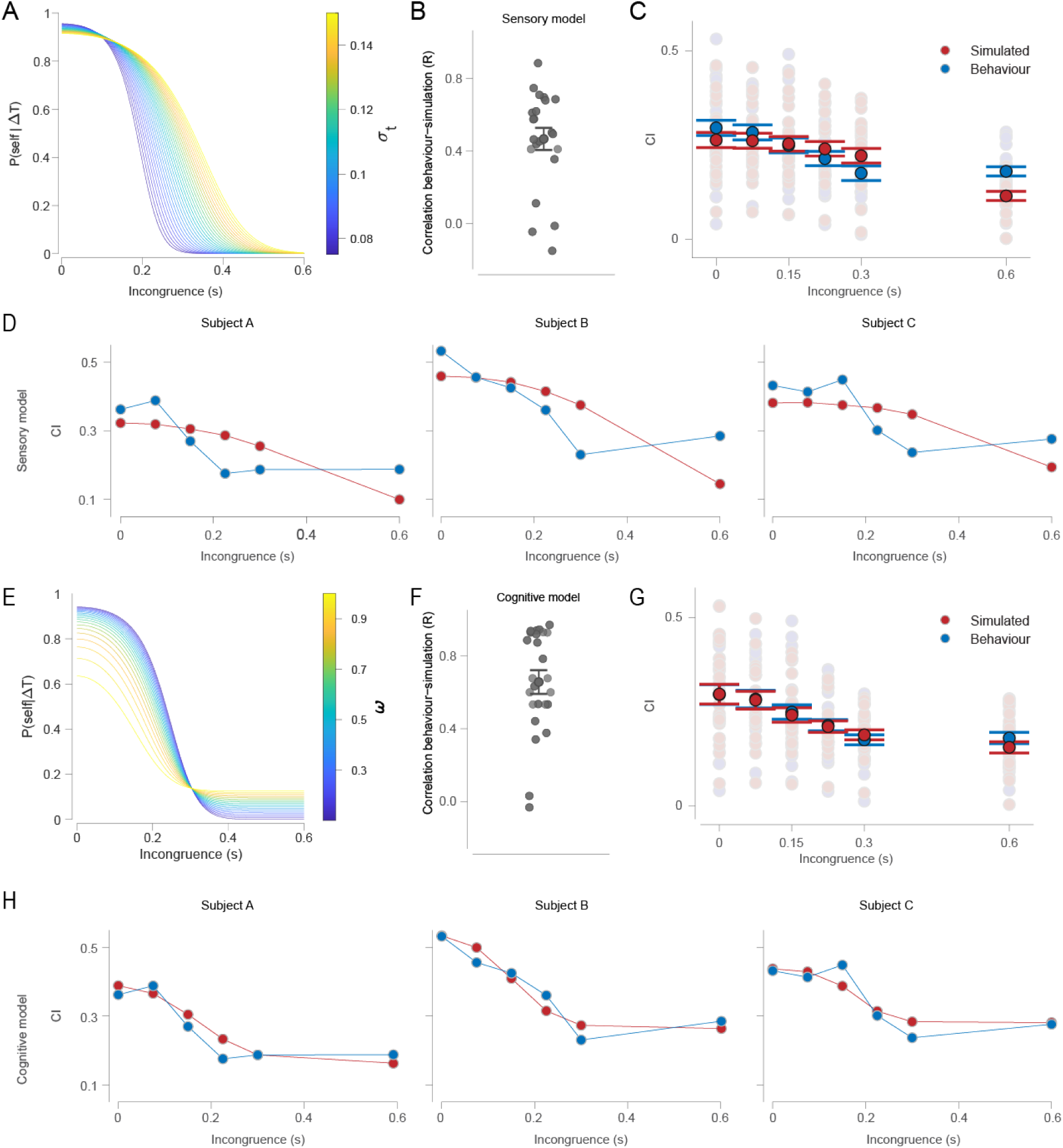
Bayesian model predictions and comparison with behavioural data. (A) Effect of temporal precision σ_t_ on P(self | Δt) in the sensory model. Each line represents model estimates for P(self | Δt) as a function of temporal incongruence, for a different value of σ_t_, chosen within a plausible range (0.075 to 0.15 s). Each other parameter was kept fixed at plausible values. (B) Correlation values between predicted and empirical CI values for the sensory model for each participant. Error bars represent standard errors. (C) Comparison between simulated (red) and empirical (blue) CI values at the population level for the sensory model. Error bars represent standard errors. (D) Sensory model predictions and empirical CI values for three exemplary subjects where the sensory model fails to capture the saturation of CI at non 0 values. (E) Effect of the cognitive parameter ω on P(self | Δt) in the cognitive model. Each line represents model estimates for P(self) as a function of temporal incongruence, for a different value of the cognitive term ω. For increasing values of ω, P(self | Δt) tends to saturate at increasing non 0 values. (F) Correlation values between predicted and empirical CI values for the cognitive model for each participant. Error bars represent standard errors. (G) Comparison between simulated (red) and empirical (blue) CI values at the population level for the cognitive model. Error bars represent standard errors. (H) Cognitive model predictions and empirical CI values for the same three subjects as in panel (D).

The model performed well overall, with an average correlation value of 0.47 ± 0.06 SEM between empirical and simulated data (Figure 4B). However, at a closer inspection, we found that the model did not fully capture the sigmoidal shape of the measured behaviour, as qualitatively apparent in Figure 4C. To explore the cause of this phenomenon we evaluated the fits at the participant level and observed that the empirical CI sigmoids saturated at non zero values for most participants (Figure 4D). This was also the case for agency ratings. The reason our model is unable to reproduce such behaviour is because *P*(*self* | *Δt*) inevitably saturates to 0 at long temporal delays for reasonable values of the temporal precision *σ_t_*. One possible explanation for the saturation of empirical CI at non 0 values is that the inference is not purely based on bottom-up sensorimotor congruence, but also on contextual cues integrated over longer timescales. These could lead to deduce visual feedback is self-caused even in the presence of very long delays. We thus added a term to Equation 3 to include such contextual inference in our model. This “cognitive term” accounts for the possibility that self-generated feedback may be presented with a longer temporal delay than what is expected purely based on internal neural noise, translating the fact that participants may be aware that a delay could have been artificially introduced (Equation 4):

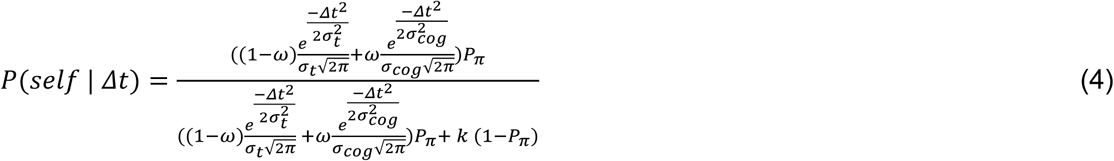

Where *σ_cocc_* is the (fixed) expected variability of artificial delay and the cognitive parameter *ω* is the weight of contextual cues in the inference. Qualitatively, this model allows for *P*(*self* | *Δt*) to saturate at non 0 values for long temporal delays, with higher saturation values as the cognitive parameter *ω* increases (Figure 4E). We repeated our analyses for the model that included the cognitive term (cognitive model). The average correlation value was higher than for the first (sensory) model (0.66 ± 0.07, t(20) = 3.55, p = 0.002, Figure 4F). Qualitatively, the cognitive model better captured the behaviour both for the population (Figure 4G) and the individual participants (Figure 4H). To evaluate such differences while accounting for the increased complexity due to the additional free parameter *ω* in the cognitive model, we compared the two models using the Akaike Information Criterion (AIC). We found that the cognitive model outperformed the sensory model in 16 out of 21 participants, showing a lower AIC value at the group level (average AIC difference = 4.64 ± 1.46, t(20) = 3.2, p = 0.0045, see also Figure S5).

## Discussion

Through a combined behavioural-computational approach, we tested the hypothesis that sense of agency is the subjective counterpart of feature selection for closed-loop motor control. To do so, we developed a task allowing us to manipulate and measure the sense of agency, while independently measuring the amount of visuomotor integration. If our hypothesis is valid, then correctable perturbations to visual feedback should only enter the closed-loop (and therefore be compensated for) when participants feel like they are causing visual feedback. Indeed, we found that the compensation index (CI), our measure of visuomotor adaptation, and explicit agency ratings were similarly influenced by temporal incongruence (Figure 1C-D). Qualitatively, when incongruence exceeded 0.175 seconds (consistent with previous reports, e.g., Farrer 2013), participants reported a substantial decrease in their sense of agency and showed less integration of visual feedback in their motor behaviour. Thus, when visual feedback is congruent with motor commands, it is selected for closed-loop visuomotor integration, and gives rise to a sense of agency. A link between visuomotor integration and sense of agency had been hypothesized previously (Reichenbach 2014), but such idea had not been integrated in a functional model allowing for empirical testing. By providing a parametric measure of the amount of visuomotor integration, our task allowed us to directly correlate subjective agency ratings and visuomotor integration, and provided a functionally relevant proxy of sense of agency. Indeed, in our framework, the sense of agency emerged not merely as a subjective epiphenomenon of motor control, but as the correlate of a functional signal that gated which sensory information entered the closed-loop.

Importantly, in a separate task, we measured reaction times to changes in cursor appearance as participants performed our task, as a proxy of visual attention. Incongruence still modulated visuomotor integration and sense of agency, but it did not affect participants’ reaction times (Figure 3C). Furthermore, there was no correlation between agency ratings or CI and reaction times. In line with previous reports (Reichenbach 2014), this suggests visuomotor binding and the associated experience of agency rely on a different mechanism than visual attention.

To further assess the link between visuomotor integration and sense of agency, we correlated variability in agency ratings with visuomotor adaptation at the single trial level, after regressing out the effect of incongruence. As expected, most variability in both measures was explained by temporal incongruence (Figure 2B), but agency ratings still correlated significantly with the amount of visuomotor integration when temporal incongruence was factored out (Figure 2C). This correlation is especially significant because it implies covariation at the single trial level extending beyond the effect of delay. This rules out that agency and visuomotor integration may be simply unrelated by-products of temporal incongruence, and suggests that the same neural process may lead to visuomotor integration and sense of agency. We nonetheless refrain from establishing a full equivalence between closed loop visuomotor integration and all aspects of the control experience. This is especially important when accounting for the biases and limitations of explicit reports. Indeed, subjective agency reports capture a complex and multi-faceted phenomenon, reflecting not only the overall self-causation feeling (that is, sense of agency in the strict sense), but likely also the experienced smoothness and precision of the control. Only the former should drive feature selection in an optimal closed-loop controller, and thus be reflected in visuomotor integration. Indeed, small levels of incongruence may affect the perceived smoothness of the control experience, but not the overall tendency to interpret feedback as self-caused, and thus integrate and correct errors.

These subtleties highlight the value of formalising this process in a rigorous modelling framework, in which distinct components can be explicitly defined and quantitatively related. We modelled behaviour in our task as the result of a Bayesian inference process, whereby motor output reflects the weighted combination of visual and proprioceptive signals. The relative weight attributed to visual information (and thus the CI) is proportional to the probability that its movement is self-caused, which is in turn inferred based on temporal congruence between hand movements and cursor movements. A purely bottom-up model, assuming that only minimal delays are compatible with self-causation, was outperformed by a model allowing longer delays to remain consistent with self-causation (Figure 4F-G). The latter captured the contribution of higher-level, temporally extended inference processes that cognitively integrate contextual information. This is in line with modern accounts suggesting that both top-down and bottom-up factors contribute to the sense of agency (Moore and Fletcher 2012). Importantly, from the computational standpoint, the output of the process is simply a self-causation probability, containing no information about the top-down or bottom-up nature of the inference. The associated phenomenology, however, is likely to be more complex.

Beyond the behavioural and computational levels, an important path for future research will be to focus on the neural correlates of the different components of self-causation inference and the associated motor control loops. It is tempting to speculate that motor control associated with bottom-up inference may be implemented by visuomotor loops in frontoparietal networks (Desmurget et al. 2009; Blakemore and Sirigu 2003), while control linked top-down inference may involve more frontal areas involved in attentional control and executive function (Synofzik et al. 2009). While we observed that both processes can lead to some extent of closed-loop integration, bottom-up control yielded much higher CI values, and is likely associated with a natural, effortless control acting on faster timescales. Understanding the behavioural and neural principles of these different modes of control may help develop more effective neuroprostheses and brain-machine interfaces, and accurately monitor user experience.

While we used visuomotor integration as a study case, our framework also directly applies to proprioceptive feedback. In able-bodied individuals, proprioceptive feedback is normally congruent with motor commands, and thus should by default be integrated with them according to our framework. However, when sensorimotor loops are altered (e.g., spinal cord injury, stroke, amputation), the integration between proprioceptive feedback and motor commands should depend on the variable degree of perceived congruence between the two. Importantly, our modelling framework predicts that the relative weights of proprioceptive and visual signals in sensorimotor integration will depend on their reliability. This can be tested and applied in neuroprosthetics. Indeed, embodiment and agency are key aspects in the success of neuroprosthetic devices (Maimon-Mor and Makin 2020), but methods to measure it in this context suffer from limitations, especially explicit questionnaires (Zbinden et al. 2022; Segil et al. 2022). Our implicit approach can be applied to extract objective behavioural measures of agency, allowing to trace and compare subtle changes in the control feeling throughout and across rehabilitation protocols. Importantly, our measure avoids the biases and demand characteristics of subjective reports, making it especially suited for repeated measurements, typically required to follow neurorehabilitation trajectories. Our implicit measure could also be especially useful to reliably detect alterations in sensorimotor integration and sense of agency in neuropsychiatric conditions such as autism and schizophrenia (Gallagher and Trigg 2016; Zalla and Sperduti 2015), where questionnaire biases can be especially limiting.

In sum, we showed that closed-loop integration is closely linked to the subjective experience of agency. We provided a behavioural framework allowing us to quantify both these aspects of motor control, and a computational framework modelling them as the result of a combination of bottom-up and top-down inference of self-causation. This framework may be especially useful to study the neural correlates of such processes, and provide valuable insights in neuroprosthetics and in neuropsychiatric conditions.

## Methods

### Main behavioural task and participants

We recruited 21 right-handed, healthy participants (4 female, 17 male, mean age: 28.4) naive to the task. The experiments were conducted on a Lenovo ThinkPad E14 Gen6 laptop with a screen refresh rate of 60 Hz. All experiments were approved by the Geneva canton ethical board (CCER) and participants were compensated 20.– CHF/hour. Participants were instructed to perform circular movements with a manipulandum (Thrustmaster T.Flightstick X, sampled at 20 ± 3 Hz) whilst looking at a fixation cross. They were instructed to keep a steady pace (around 1.5 circles per second) and pay attention to the cursor. To isolate the effect of subconscious adaptation, they were explicitly advised against slowing down to correct their movements or trying to “draw” circles on the screen with the cursor. Participants pressed a button on the manipulandum to start the trial, followed by a one second preparatory cue, followed by nine seconds of motor activity. After each trial, participants reported their subjective experience of agency for the cursor on a 1-9 visual analog scale. Every participant completed a total of 248 trials: 5 distortions by 6 incongruences, plus one open loop (no visual feedback) condition, each condition being repeated 8 times, in a completely randomized order. The task was divided in five blocks lasting about 12 minutes, for a total duration of about 75 minutes including breaks. Sample size was determined heuristically based on pilot data from 5 participants, who all showed a decreasing trend of CI and agency with incongruence. We thus aimed at obtaining data from 20 participants, expecting this would be largely sufficient to reach statistical significance in these main analyses. 21 participants were recruited to protect against data loss and technical issues, and none had to be discarded in the end, yielding the final sample size of 21 participants.

### Temporal incongruence and spatial distortions

The temporal incongruence was implemented by adding a non-constant delay to the displayed location of the effector. For each incongruence, delays were sampled from Gaussian distributions with mean and standard deviation respectively: [(0, 0.1) (0.075, 0.15) (0.15, 0.3) (0.225, 0.45) (0.3, 0.6) (0.6, 1.2)]. Delays were clamped to non-negative values, meaning they were set to zero if a negative value resulted from Gaussian sampling. A trial started with the sampling of a delay. During the delay period, visual feedback was displayed with the corresponding delay. Once a delay ended a new delay value was sampled from the distribution. In order to present a continuous trajectory of the dot when a new delay was sampled, the trajectory was smoothed by displaying the exponential moving average of the cursor, with a smoothing factor that depended on movement speed, so faster movements were smoothed more. The correctable distortions were implemented as a multiplicative factor (0.7, 1, 1.3) of the mapping of the input on either the horizontal or vertical axes. For simplicity in analysing the data and presenting our results, we projected all distortions to the horizontal axis, so that vertical distortions got converted to the following values: 1/1.3 = 0.769, 1, 1/0.7 = 1.429. All code for the task was implemented in python.

### Reaction time task

We had the same 21 participants complete a variation of our main task studying visual attention. They followed the same instructions as in the main task, but they were also instructed to press a trigger on the manipulandum as quickly as possible if the cursor changed colour. Distortions were the same as in the main task, but only two temporal incongruences were presented [(0, 0.1) (0.6, 1.2), mean and SD]. For each condition, the cursor changed colour in half the trials to prompt a reaction. Every participant completed 120 trials: two incongruences, three distortions and reaction or non-reaction type trials, each repeated 10 times in randomized order.

### Behavioural data analysis

Preprocessing of the data was performed in Python. First, the time series data of each trial was segmented in individual circular movements (segments) by dividing the manipulandum square range of motion in four quadrants. A segment was defined whenever the manipulandum visited each of the four quadrants consecutively. Ellipses were fitted to each segment, and the associated metrics were returned for further analyses implemented in R. The ratio between the ‘width’ and ‘length’ from the elliptical fit (from now: elongation) was computed as our metrics of interest. Elongations from the first three segments were discarded to remove the transient adaptation period from our analyses (see Figure S1). Furthermore, outliers deriving from noisy or non-elliptical trajectories leading to poor elliptical fits were filtered out by evaluating the median absolute deviation of the elongation. A segment was rejected if it had a median absolute deviation above 2 compared to all segments in the current trial (mean rejection rate of 9.1%). Then, the median elongation across segments for each trial was computed and used to extract the compensation index for each temporal incongruence. To do this, we ran a linear regression predicting elongation values from distortion values for each incongruence, and extracted the negative slope of such regression. This provides a condensed index of the amount of motor adaptation for a single incongruence, the compensation index (CI). A CI of 0 indicates that motor behaviour is fully independent from spatial distortions. A CI of 1 indicates that motor behaviour fully adapts to spatial distortions, so that the trajectory of the cursor is independent from spatial distortions.

### Residual analysis

To calculate the residuals for agency ratings we subtracted to each agency rating the mean agency rating computed over each incongruence and participant. Because the CI is calculated through a regression over a multitude of trials, to obtain a single trial proxy of visuomotor integration, we calculated the behavioural residuals by evaluating the relative elongation in a single trial. This was achieved by subtracting to each elongation the average elongation computed over each incongruence and participant. Residuals corresponding to distortions below 1 were flipped in sign, as more negative values translate to more compensation relative to the mean elongation in this case. By this, we achieved values that increase with the amount of elongation irrespective of the direction of the distortion. Trials with no distortion were excluded from the analysis, as they reflect individual biases or noise and provide no meaningful information about adaptation. After these steps, a positive residual elongation indicated more integration of visual information.

### Bayesian Model of Behaviour

The core principle of our model is that the inference of self-causation influences motor behaviour by affecting the perceived sensory consequences of actions, and thus the adaptation response towards the desired behaviour. Thus, our model focuses on perceptual aspects, assuming that motor behaviour will adapt to obtain what is perceived as the desired behaviour. We assume that, in normal conditions, proprioceptive feedback is always perceived as self-caused. In case visual feedback is also perceived as self-caused, it will be subconsciously integrated with proprioceptive feedback to form a unified percept about the outcome of motor commands. Thus, even if participants are instructed to perform circles with their real hand and not with visual feedback, they are expected to correct for distortions of visual feedback. In case visual feedback is not perceived as self-caused, participants will only adjust behaviour based on proprioceptive feedback.

As routinely done in Bayesian models of behaviour (Rohe and Noppeney 2015), we will start by defining the generative model of sensory inputs, and then apply Bayes’ theorem on it. We model the sensory feedback encoded in the visual and proprioceptive modality (X_*vis*_, X_*prop*_) as the sum of the actual sensory stimulus and neural noise.

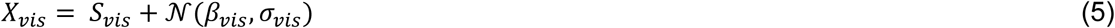

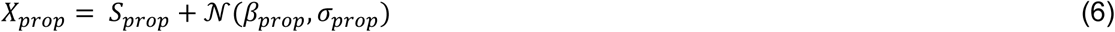

Where *S_vis_* and *S_prop_* represent the true elongation of visual and proprioceptive elliptical trajectories, *β_vis_* is and *β_prop_* represent a fixed perceptual bias, *σ_vis_* and *σ_prop_* represent the encoding noise in the visual and proprioceptive modalities. X*_vis_*, X*_prop_* thus represent the encoded elongation of visual and proprioceptive trajectories respectively.

Similarly, the encoded temporal incongruence is modelled as the true incongruence plus a Gaussian neural noise term.

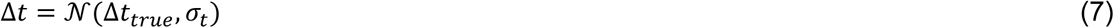

In the case of non-self-causation, the probability of observing a delay *P*(Δ*t* | ! *self*) is modelled as a constant *k*, fixed at 1 for simplicity.

The probability of observing a delay in the case of self-causation determines the difference between the sensory and the cognitive model. In the sensory only model, true delays are assumed to be always zero, and the distribution of observed delays is simply a Gaussian distribution centred at zero, with variability determined by neural noise *σ_t_*.

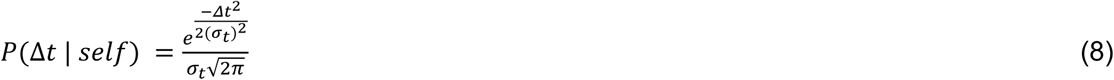

The cognitive model accommodates for the possibility that true delays may be artificially introduced even in the case of self-causation. Thus, the distribution of delays consists of the mixture of two gaussians. The first gaussian is the one of the sensory model (sensory noise, no artificial delay). The second gaussian, with variance fixed at 10 seconds, is meant to approximate a uniform distribution on a broad range of values, representing the possibility that a delay is artificially induced. The relative weight of the two gaussians is governed by the parameter *ω*, representing the expected probability that a delay could be artificially induced.

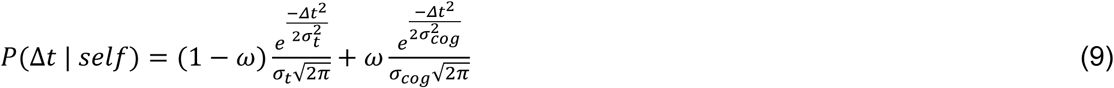

We assume that the inference of self-causation is only dependent on this observed temporal delay, and apply Bayes’ theorem to compute the inferred probability of self-causation.

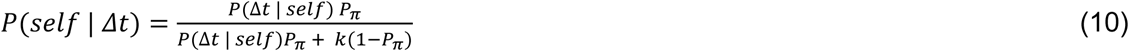

Where *Pπ* is the individual prior for self-causation.

After deriving the expression for self-causation probability, we model its effect on behaviour. As in previous similar models (Bertoni et al. 2023; Fang et al. 2019), we assume that the final perceived trajectory will be the optimal integration of visual and proprioceptive inputs (as in Ernst and Banks, 2002) weighted by the self-causation probability *P*(*self* | *Δt*), plus proprioceptive inputs only weighted by 1 − *P*(*self* | *Δt*):

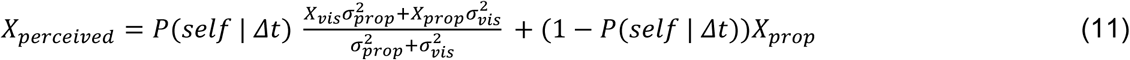

The motor behaviour *Ŝ*, is finally modelled with a single shot correction of X*_perceived_*. In other words, we assume subjects to adapt their motor behaviour so that it would lead X*_perceived_* to be 1 (a circle), as per our instructions. Gaussian noise of variance *σ_out_* is added to account for noise in the implementation of the motor plan.

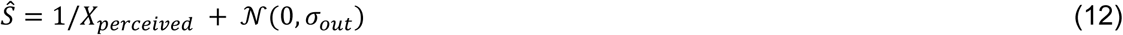

Finally, we model the effect of distortions in our task by simply introducing a relation between true visual and proprioceptive inputs:

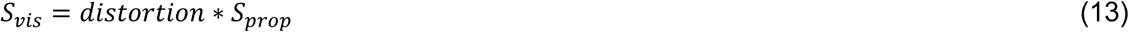

Temporal incongruences do not, on average, affect the relation between *Svis* and *Sprop*.

### Model fitting

The sensory and cognitive models have 7 or 8 free parameters respectively: *σ_vis_*, *σ_prop_*, *σ_t_*, *σ_out_*, *β_vis_*, *β_prop_*, *P_π_* and *ω* for the cognitive model only (*σ_cocc_* is fixed). These parameters were fitted individually to each participant by simulating 10000 samples of behaviour for each distortion and incongruence, and then finding the set of parameters that maximize the likelihood of the data given the simulated model predictions through the MATLAB function ksdensity, similarly to what was done in (Bertoni et al., 2023). Parameters were optimized through stochastic gradient descent implemented in MATLAB using the BADS toolbox (https://github.com/lacerbi/bads) (Acerbi and Ma, 2017). Fits were repeated 10 times for each subject with different starting parameters, and the iteration yielding the highest log-likelihood was selected. Open-loop trials were included in model fitting to help the estimation of motor noise and individual biases. An example of model fitting is shown in Figure S6, fitted parameters for each model are shown in Figure S7.

### Model evaluation

To evaluate model performance, for the set of parameters extracted for each participant, we simulated 10000 trials for each incongruence, distributed on the same distortions as in our behavioural task. CI values were extracted from simulated behaviour and compared to empirical CI values by comparing the correlation coefficient. To compare the two models while accounting for the greater complexity of the cognitive model, we used the Akaike Information Criterion (AIC): AIC=2k-2ln(L), where k is the number of model parameters, and L is the likelihood of the best fitting model. Log-likelihood values were directly computed by the fitting algorithm as described above.

## Notes

### Competing Interest Statement

The authors have declared no competing interest.

### Funding

- Swiss National Science Foundation: 223402 Ambizione grant awarded to Tommaso Bertoni

