## Supplementary Information for "A Unified Neurocomputational Framework for Closed-Loop Motor Control and Sense of Agency"

**Supplementary Figures**

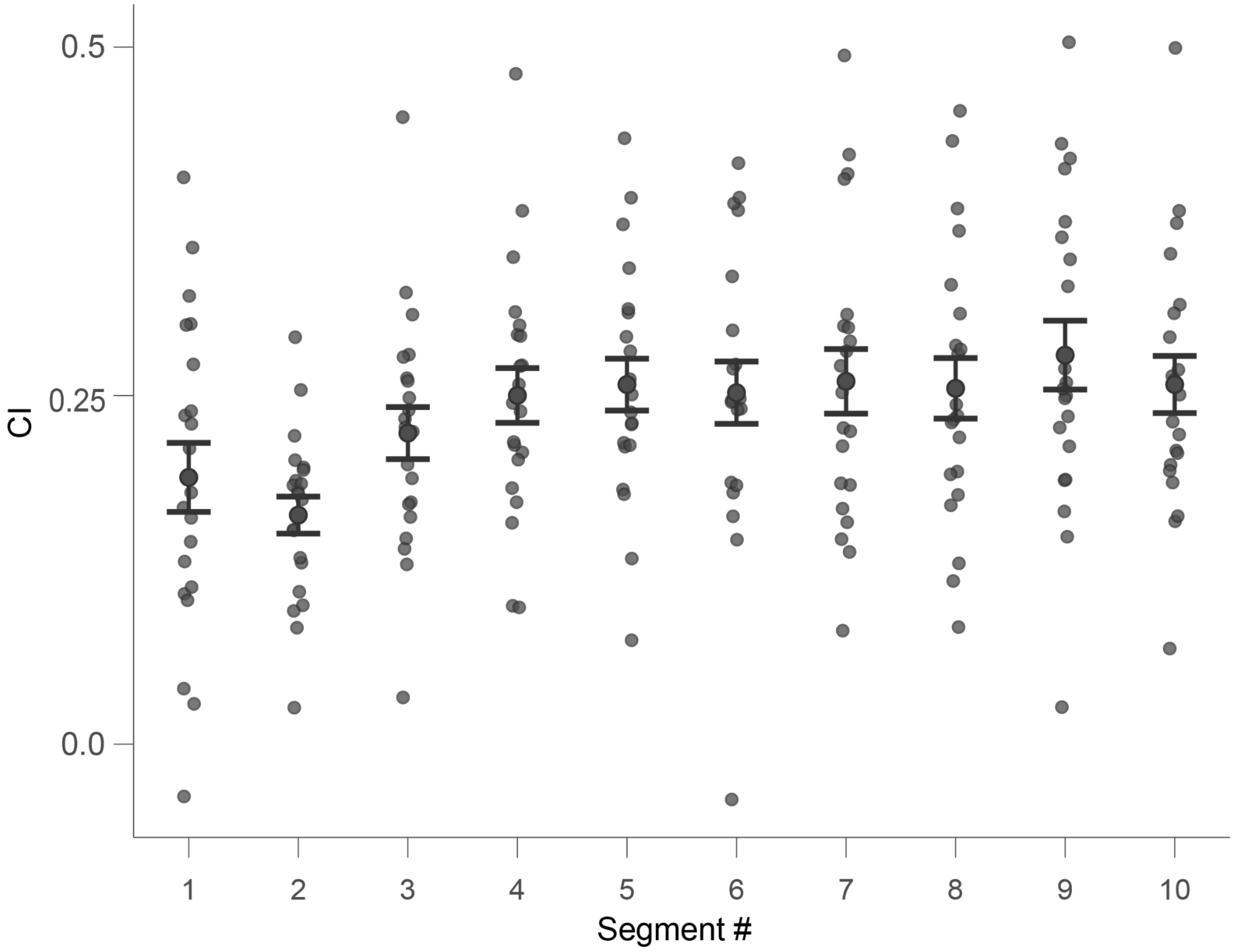


***Figure S1.*** *Compensation indexes across individual segments. To remove the transient adaptation time at the beginning of each trial, we studied the temporal dynamics of the CI (averaged across incongruences) across individual segments. Since all subjects completed at least 10 segments on each trial, we focused our analysis on the first 10 segments of each trial. Visually, the CI appears to stabilise to a fixed value starting from the fourth segment. To confirm this, we compared CI values for each segment to the saturation value defined as the average across the last four segments. The first three segments were significantly different from the saturation value ($p = 0.009$, $p < 0.0001$, $p = 0.045$ respectively), while the remaining segments were not (all $p > 0.52$).*

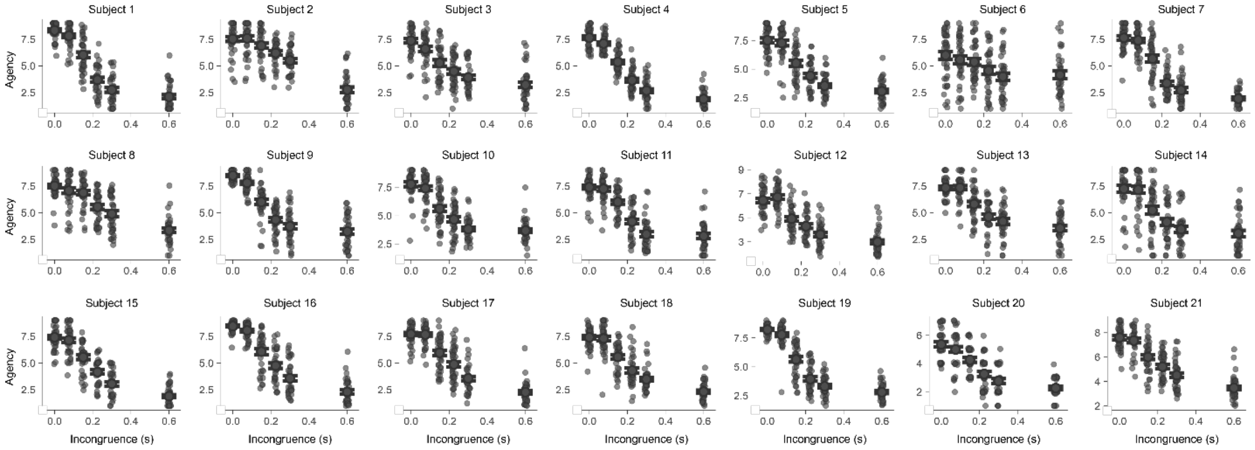


***Figure S2.*** *Agency ratings for every trial plotted against the incongruence for each participant. Errorbars indicate standard errors.*

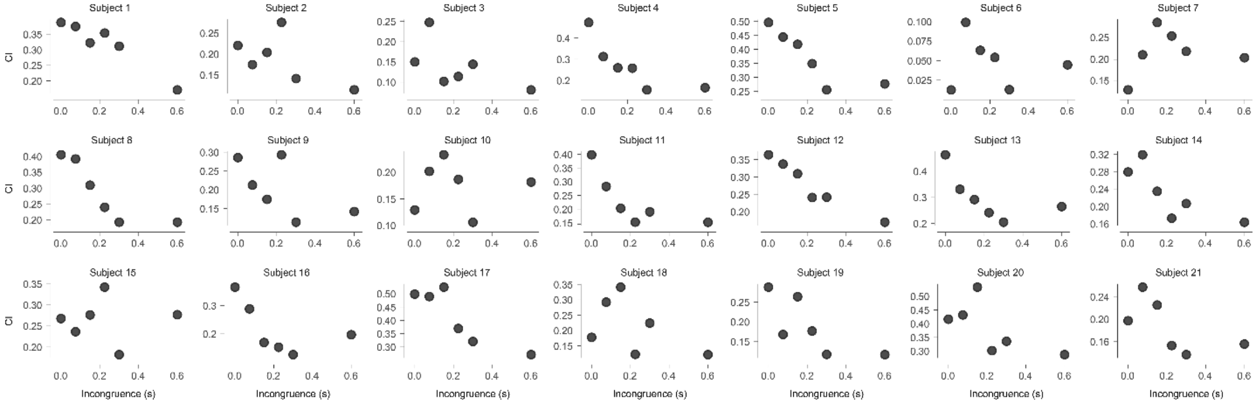


***Figure S3.*** *CI for each incongruence and participant.*

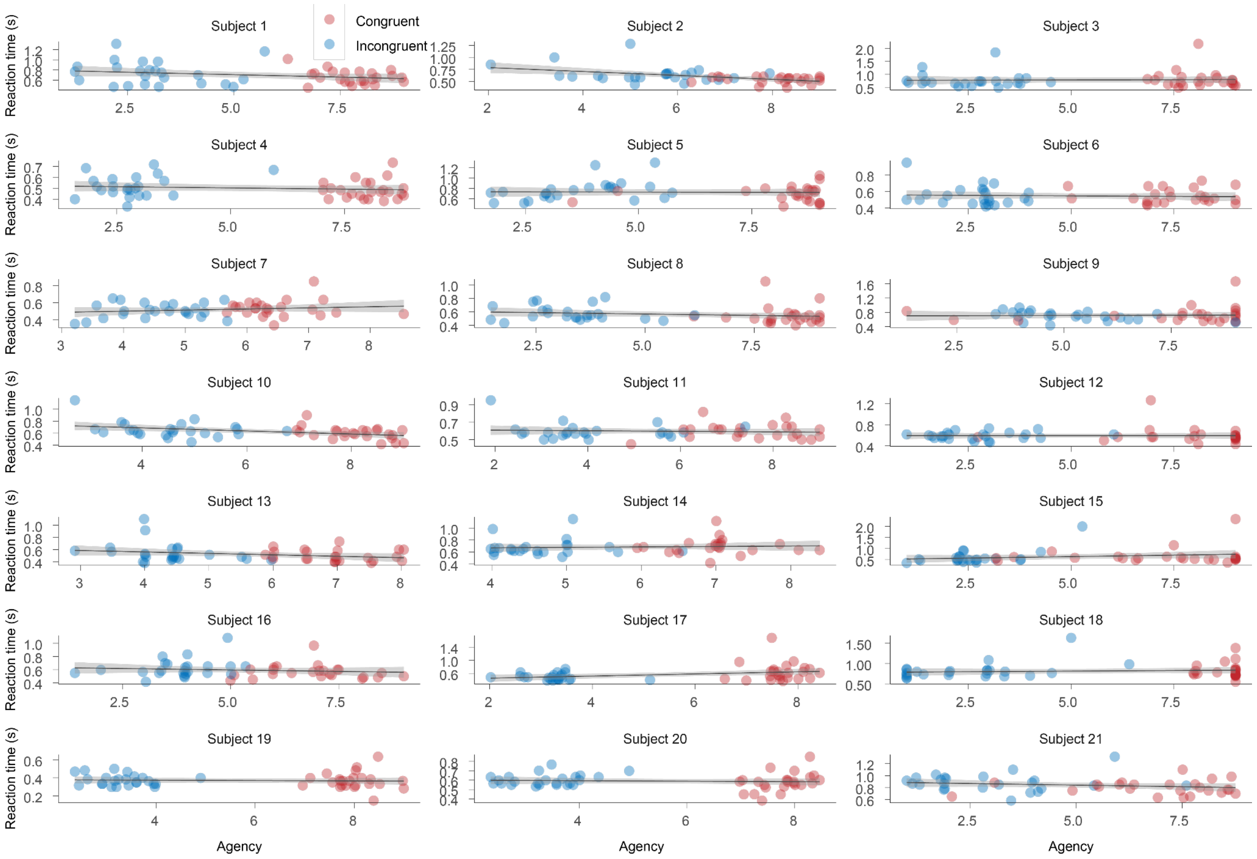

***Figure S4.*** *Linear fit of reaction times and agency ratings per participant, colour indicates congruence. Only three of the subjects have a significant linear fit (subject 2: R=-0.482 p<0.001, subject 10: r= -0.41 p=0.004, subject 17: r=0.336, p=0.02).*

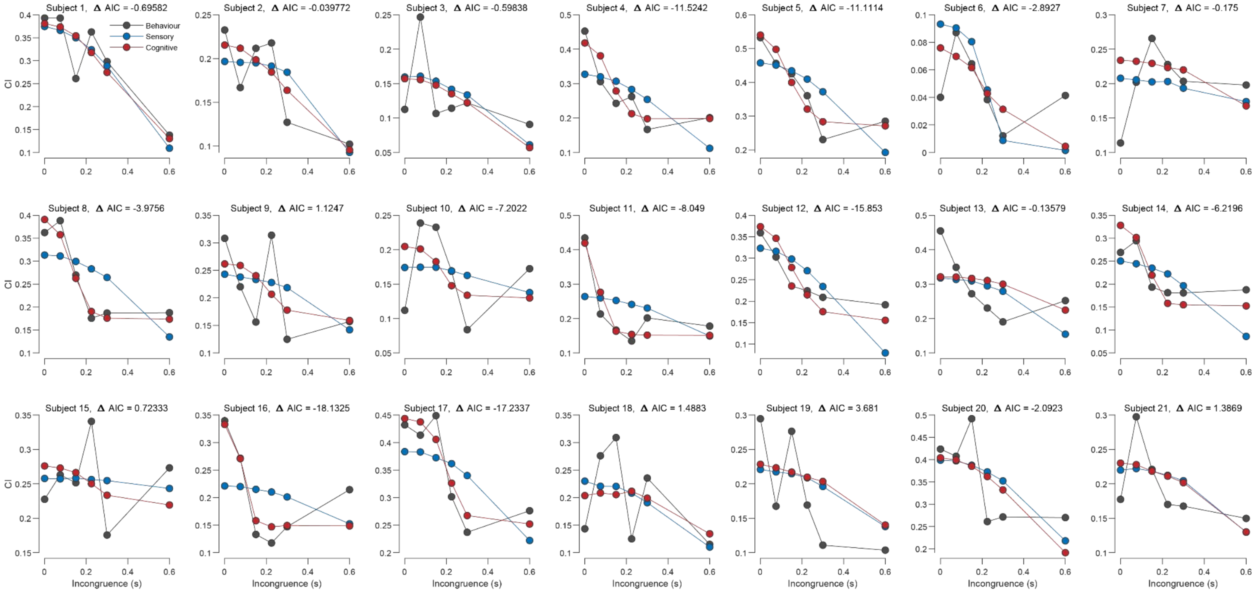

***Figure S5.*** *Comparison of sensory and cognitive model at the single subject level. Negative Δ AIC values favour the cognitive model.*

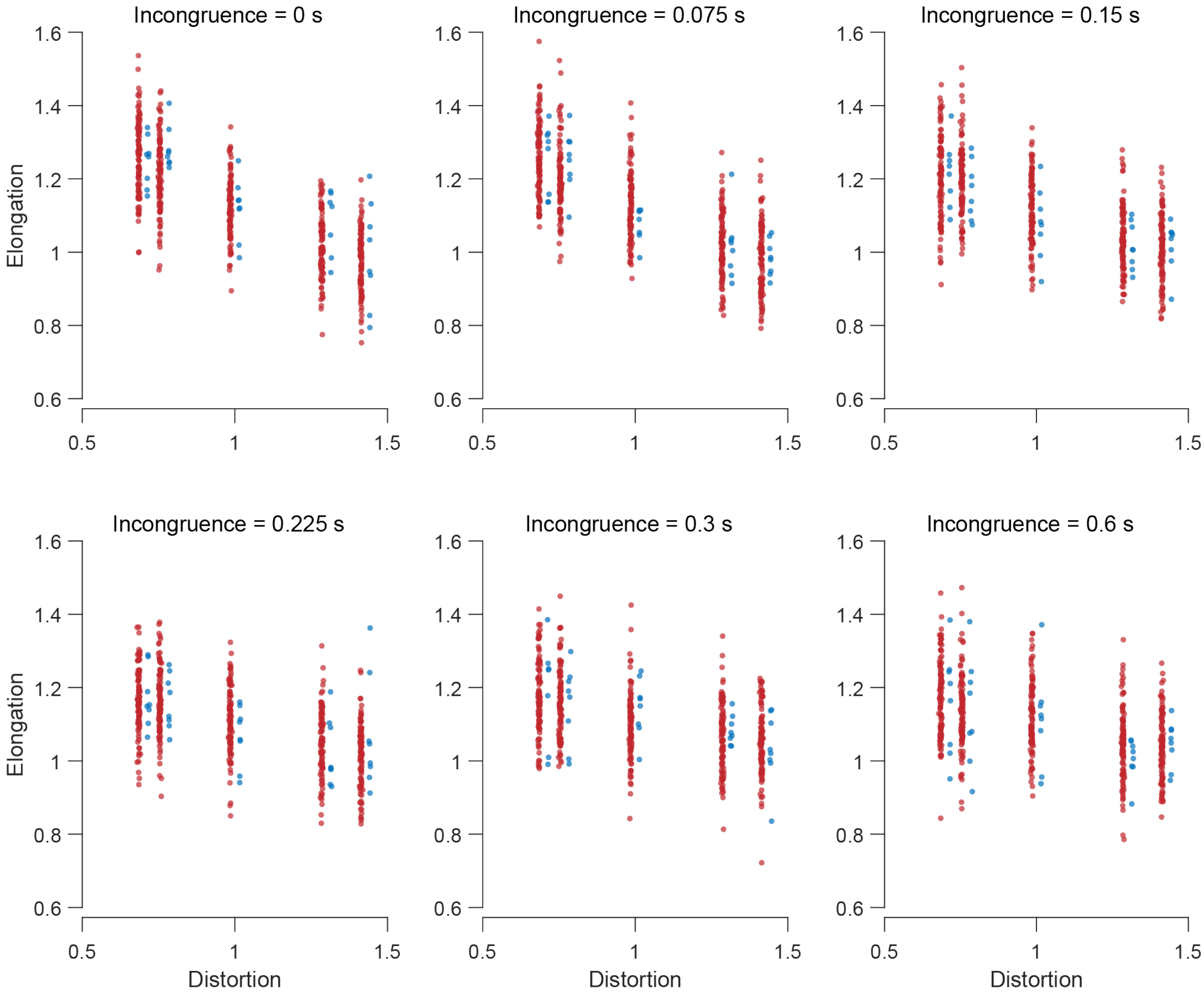


***Figure S6.*** *Model fitting for an example participant. Each panel indicates a temporal incongruence. Blue dots indicate elongation values for individual trials. Red dots indicate 100 simulated data points for best fitting parameters for this participant. Model fitting was performed by simulating 10000 trials per incongruence and distortion.*

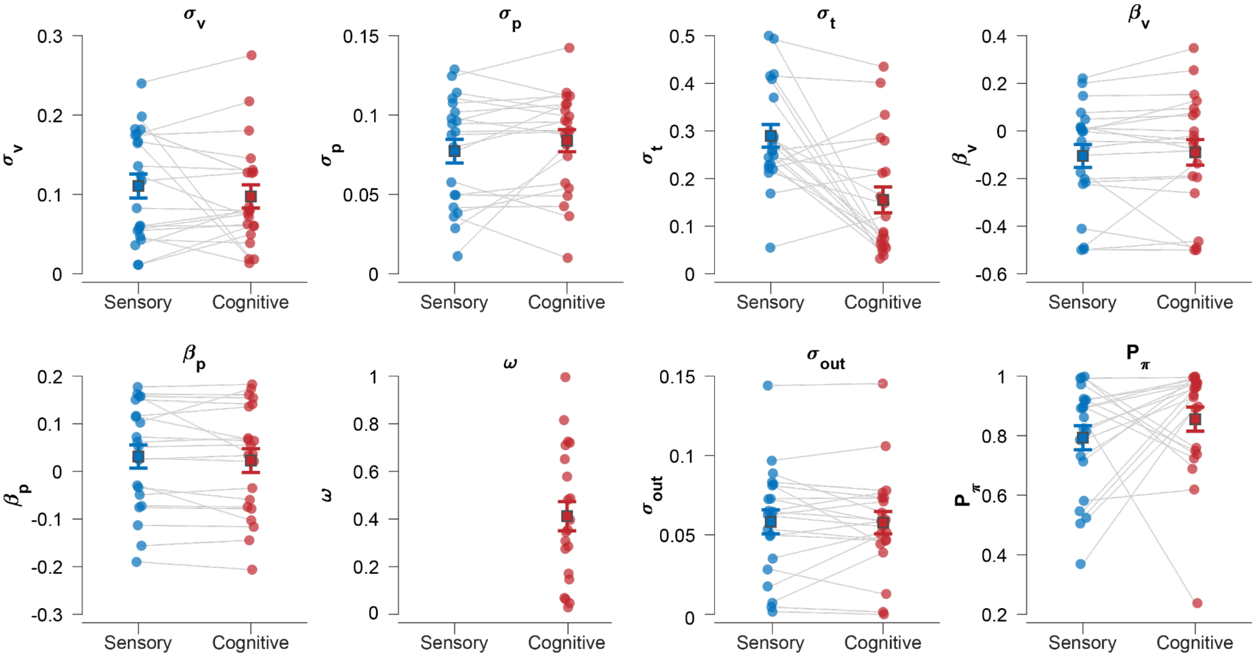


***Figure S7.*** *Fitted parameters for the sensory and cognitive model. Error bars represent standard errors. Most parameters are relatively stable across the two models, with the notable exception of the temporal precision $\sigma_t$, which is lower in the cognitive noise. This is likely because the sensory model attempts to account for non-zero saturation values by increasing values of $\sigma_t$.*
